# Genome-wide Identification of Zero Nucleotide Recursive Splicing in *Drosophila*

**DOI:** 10.1101/006163

**Authors:** Michael Duff, Sara Olson, Xintao Wei, Ahmad Osman, Alex Plocik, Mohan Bolisetty, Susan Celniker, Brenton R. Graveley

**Affiliations:** Department of Genetics and Developmental Biology, Institute for Systems Genomics, University of Connecticut Health Center, Farmington, Connecticut 06030, USA; Department of Genome Dynamics, Lawrence Berkeley National Laboratory, Berkeley, California 94720, USA.

## Abstract

Recursive splicing is a process in which large introns are removed in multiple steps by resplicing at ratchet points - 5’ splice sites recreated after splicing^1^. Recursive splicing was first identified in the *Drosophila Ultrabithorax* (*Ubx*) gene^1^ and only three additional *Drosophila* genes have since been experimentally shown to undergo recursive splicing^2,3^. Here, we identify 196 zero nucleotide exon ratchet points in 130 introns of 115 *Drosophila* genes from total RNA sequencing data generated from developmental time points, dissected tissues, and cultured cells. Recursive splicing events were identified by splice junctions that map to annotated 5’ splice sites and unannotated intronic 3’ splice sites, the presence of the sequence AG/GT at the 3’ splice site, and a 5’ to 3’ gradient of decreasing RNA-Seq read density indicative of co-transcriptional splicing. The sequential nature of recursive splicing was confirmed by identification of lariat introns generated by splicing to and from the ratchet points. We also show that recursive splicing is a constitutive process, and that the sequence and function of ratchet points are evolutionarily conserved. Together these results indicate that recursive splicing is commonly used in *Drosophila* and provides insight into the mechanisms by which some introns are removed.

---

Recursive splicing was first identified in the *Drosophila melanogaster Ultrabithorax* (*Ubx*) gene^1^. The 73 kb intron within *Ubx* houses two alternative microexons (mI and mII) which both contain the consensus 5’ splice site sequence GTAAGA immediately downstream of the 3’ splice sites. In addition, this intron contains a ratchet point, a zero nucleotide exon consisting of juxtaposed 3’ and 5’ splice sites. It has been shown that rather than being removed in a single step, the 73 kb *Ubx* intron is removed in four steps in which the upstream constitutive exon is spliced to exon mI, and subsequently re-spliced to exon mII, the ratchet point, and finally the downstream constitutive exon (Supplementary Fig. 1). A previous genome-wide computational search for potential ratchet points conserved between *D. melanogaster* and *D. pseudoobscura* predicted 160 potential ratchet points in 124 introns of 106 genes^2^. Of these, only 7 ratchet points in three genes (*kuzbanian* (*kuz)*, *outspread (osp)*, and *frizzled (fz)*) have been reported to be experimentally validated^2,3^.

We generated 10.9 billion uniquely mapped reads of rRNA-depleted, paired-end, strand-specific RNA sequence from 183 *D. melanogaster* individual RNA samples comprising 35 dissected tissue samples, 24 untreated and 11 ecdysone treated cell lines, 30 distinct developmental stages and males and females of four strains from the *D. melanogaster* Genetic Reference Panel^4^ (Supplementary Table 1). The majority of these RNA samples were the same as those we previously used to generate poly(A)+ RNA sequence data^5,6^. As the current libraries were prepared without poly(A) selection, they contain a mixture of mRNA, pre-mRNA and nascent RNA. Co-transcriptional splicing can be observed in total, nuclear, or nascent RNA-seq data by the sawtooth pattern of read density across introns in the 5’ to 3’ direction of transcription^7^ (Supplementary Fig. 1a). While visually inspecting these data on a genome browser, we noticed several large introns that lacked internal annotated exons yet possessed sawtooth patterns of read density suggestive of co-transcriptional splicing, including the introns from *Ubx*, *kuz*, *osp*, and *fz* that were previously shown to undergo recursive splicing (Supplementary Fig. 1b). We therefore hypothesized that such sawtooth patterns could be indicative of recursive splicing and performed a genome-wide search for ratchet points supported by the RNA-Seq data.

Two independent, yet complementary approaches were used to identify potential zero nucleotide exon-type ratchet points. First, we parsed the RNA-Seq alignments to identify novel splice junctions where the reads mapped to an annotated 5’ splice site and an unannotated 3’ splice site, and the genomic sequence at the 3’ splice site junction was AG/GT (Supplementary Fig. 2a). Second, we aligned the total RNA-Seq data to a database of splice junctions between annotated exons and all potential ratchet points (AG/GT sequences) in the downstream intron that did not correspond to annotated 3’ splice sites. We then identified ratchet point junctions where reads mapped without any mismatches, with at least three distinct offsets, and with an overhang of at least eight nt (Supplementary Fig. 2b). We then visually inspected each ratchet point independently identified by both methods on the genome browser, removing candidates that did not display an obvious sawtooth pattern of read density.

We identified a total of 197 ratchet points in 130 introns of 115 genes (Supplementary Table 2). Two of these ratchet points were missed by our computational approaches, but identified during the course of manual inspection on the browser, were validated and were included in the remainder of these analyses. This provides the first experimental verification of 91 of the 160 ratchet points computationally predicted by Burnette *et al.* based on comparative genomics^2^ (Supplementary Table 3). The other 106 (53.8%) of the ratchet points we identified are described here for the first time.

Most genes (100) contain only one recursively spliced intron, though 15 genes contain two. The number of ratchet points in an intron ranges from one to six (Fig. 1a). The recursively spliced introns range in size from 11,341 bp to 132,736 bp with an average size of 45,164 bp. The introns containing recursive splice sites are enriched in large introns (97% of all introns are smaller than the smallest recursive intron), not all large introns contain recursive splice sites (Fig. 1b). In fact, only 6% of introns larger than the smallest recursive intron are recursively spliced. The segments of the introns removed by recursive splicing range from 2,596 bp to 63,580 bp with an average size of 17,953 bp (+/− 9,039 bp) and median size of 16,368 bp (Fig. 1c). The *luna* gene contains an 108 kb intron with five ratchet points, such that the intron is removed in six stepwise recursive splicing events (Fig. 1d). The five ratchet points are supported by the sawtooth pattern of read density across the intron, reads that map to the exon-ratchet point splice junctions (Fig. 1d), and have been validated by RT-PCR and Sanger sequencing (Fig. 1e). In total, RT-PCR and Sanger sequencing validated 24 ratchet points from 14 genes in *Drosophila* S2 cells (Supplementary Fig. 3).

**Figure 1.**
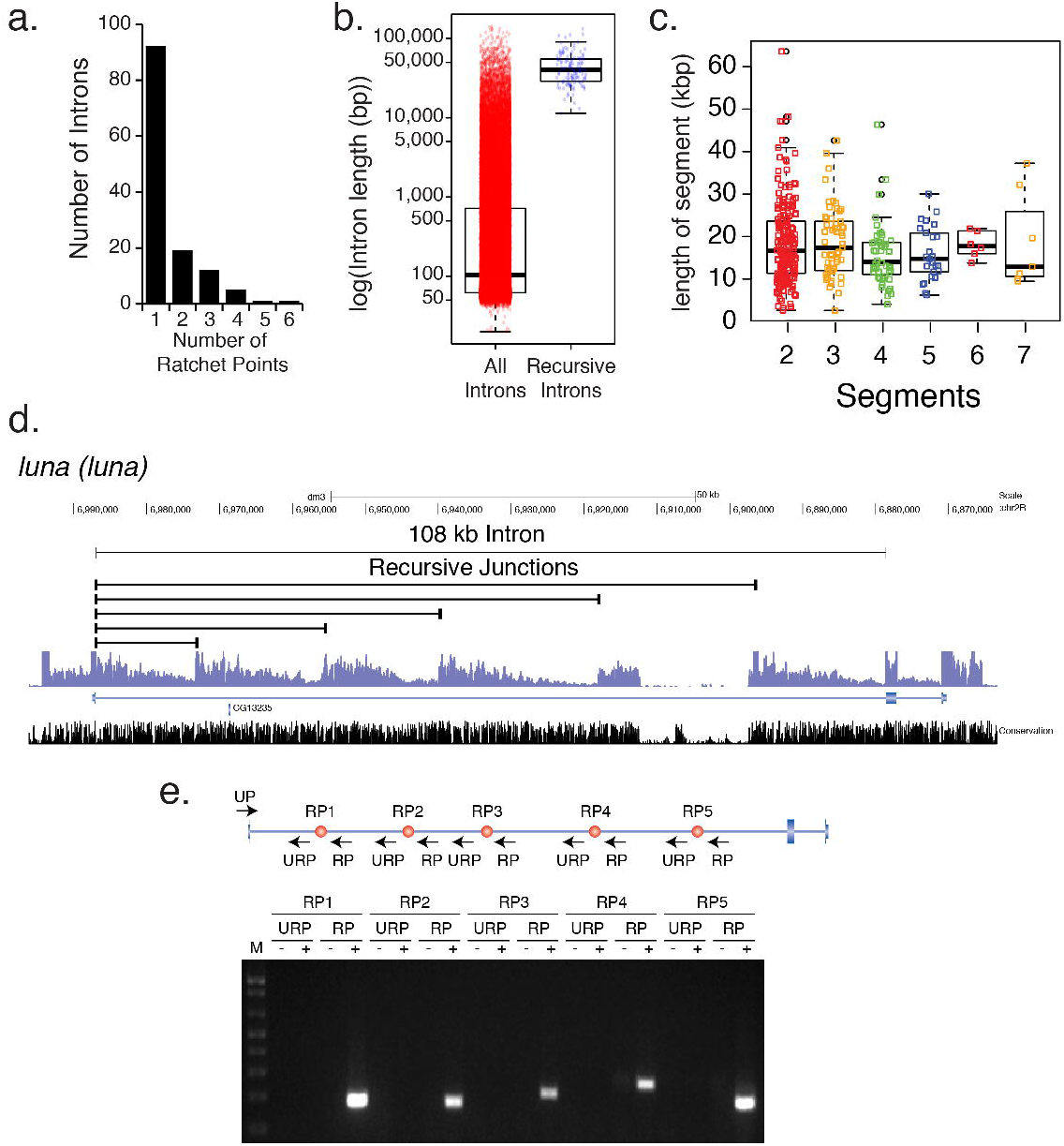
Identification and validation of new recursive splice sites. **a,** Distribution of the number of ratchet points per recursive intron. **b,** Size distribution (log_10_(bp)) of all (red) and recursive (blue) introns. **c,** Size distribution (in kbp) of the individual intron segments removed by recursive splicing binned by the number of segments per intron. **d,** Example of five recursive splice sites identified in *luna*. Shown are the recursive junctions identified, the overall RNA-Seq read density from all samples (blue), and conservation among 16 insects (black). **e,** RT-PCR validation of the *luna* ratchet points (red dots) using primers in the upstream constitutive exon and flanking the putative ratchet points (UP). The RP primers are expected to yield RT-PCR products if the constitutive exon is spliced to the ratchet point. The URP primers, which are upstream of each ratchet point, serve as negative controls.

Ratchet points are zero nucleotide exons, and therefore do not exist in the mRNA. However, direct evidence of recursive splicing can be obtained by identifying lariat introns – byproducts of all splicing reactions that contain a 2’-5’ linkage between the first nucleotide of the intron and the branchpoint. Because reverse transcriptase can occasionally traverse the branchpoint, reads corresponding to the 5’ splice site-branchpoint junction may be present in the total RNA-seq data (Fig. 2a). To identify recursive lariat introns, we generated a set of potential 5’ splice site-branchpoint junctions for all recursively spliced introns, and all possible permutations, and aligned the total RNA-seq reads to them (Supplementary Methods). Though rare, we identified 46 reads that mapped uniquely to 27 recursive lariats introns in 20 genes (Supplementary Table 4). Six of the lariat introns detected correspond to the first segment of the recursive introns and are also supported by standard splice junction reads. However, 21 of the lariat introns detected correspond to internal segments removed by recursive splicing and represent the only means of detecting their removal. For example, *couch potato* (*cpo*) contains a recursive intron with two ratchet points that is removed in three recursive splicing events, and we obtained evidence for all three lariats (Fig. 2b). This analysis also allowed us to identify the branchpoints used for these recursive splicing events. All but five of these branchpoints reside from -36 to -21 upstream of the 3’ splice site with a peak at -30 (Fig. 2b,c). Six of the 3’ splice sites appear to use two different branchpoints. We observed that 78.79% have an A at the branchpoint, while 15.15% and 6.06% have a T or C, respectively, at the branchpoint and none have a G (Fig. 2d).

**Figure 2.**
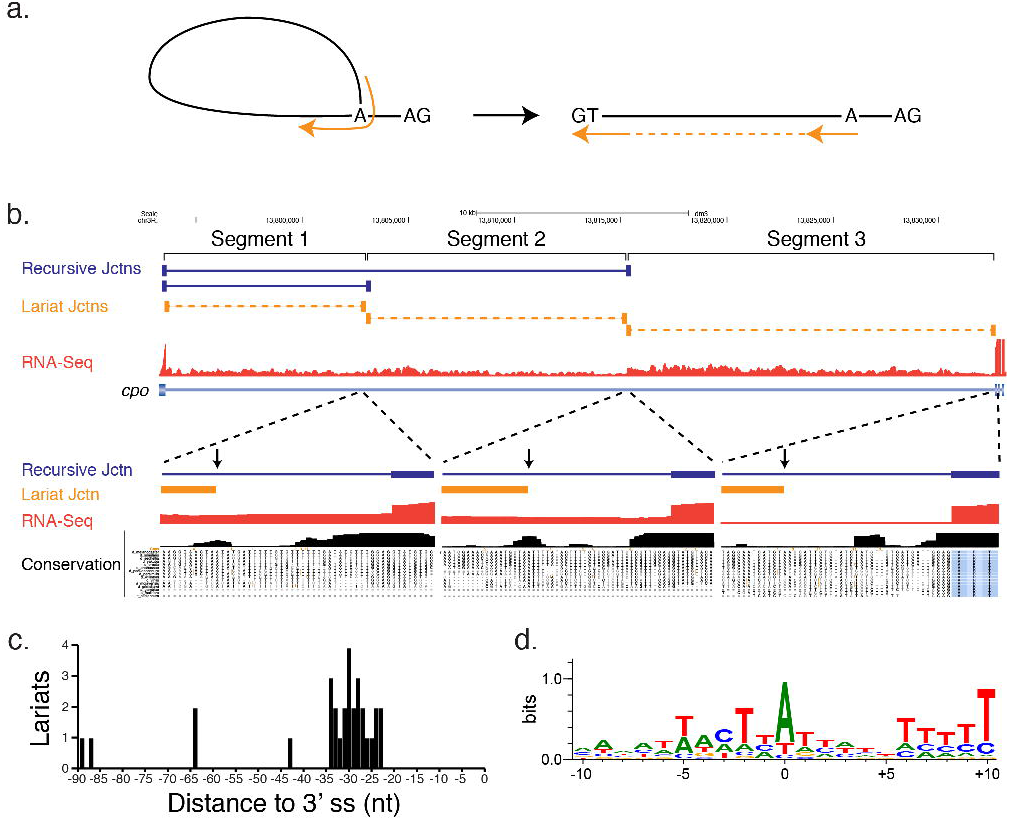
Identification of recursive lariat introns. **a,** RNA-Seq reads (orange) that traverse a 5’ splice site-branchpoint junction would aligned to the linear intron as out-of order split-reads. **b,** Example of recursive lariat introns in *cpo*. Shown are the recursive junctions identified (blue), the lariat junction reads (orange), and the overall RNA-Seq read density from all samples (red). A magnification of each branch point region is also shown along with the conservation among 16 insects. The positions of the branch points are indicated by the vertical arrows. **c,** Distribution of the distance of the recursive lariat intron branch points from the 3’ splice sites. **d,** Sequence logo of the recursive lariat intron branch point sequences.

The nucleotide sequences of ratchet points resemble juxtaposed 3’ and 5’ splice sites (Fig. 3a) and the regions immediately flanking the ratchet points are much more highly conserved than those flanking non-ratchet point AG|GT sequences in the same introns (Fig. 3b). However, the ratchet points have a more prominent pyrimidine tract, and a significantly (P=<0.0001) higher frequency of a TT dinucleotide at positions -5 and -6 relative to the 3’ splice site when compared to introns genome-wide. Whereas only 43.76% (30,151/68,898) of all introns have Ts at positions -5 and -6, 99.5% (196/197) of ratchet points do. The only ratchet point lacking a TT dinucleotide at positions -5 and -6 is in *CG15360* which has a C at position -6 that is conserved in other *Drosophila* species. Intriguingly, the majority of *C. elegans* 3’ splice sites have this sequence^8^ and it has been shown that the large U2AF subunit (encoded by *U2af50* in *Drosophila*) interacts with these bases. Thus, the strong preference for the TT dinucleotide at positions -5 and -6 of *Drosophila* ratchet points could represent high affinity U2AF binding sites so that the ratchet points are efficiently recognized.

**Figure 3.**
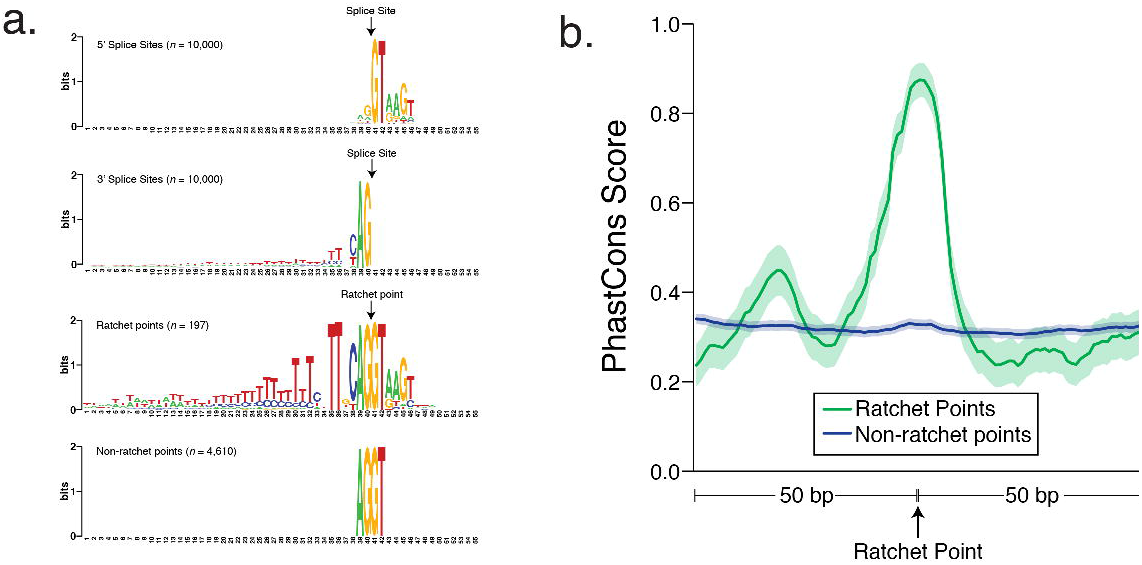
Characteristics of ratchet points. **a,** Sequence logos of 5’ splice sites, 3’ splice sites, ratchet points, and non-ratchet point AG/GT sequences located in the same introns as ratchet points (top to bottom). **b,** Sequence conservation of ratchet points. Average PhastCons scores of ratchet points (green) and non-ratchet points (blue). Solid line indicates the average PhastCons score, shaded regions indicate the 95% confidence interval.

To determine if recursive splicing is evolutionarily conserved, we generated rRNA-depleted, stranded RNA-Seq data from mixed *D. simulans*, *D. sechellia*, *D. yakuba*, *D. pseudoobscura*, and *D. virilis* adults (Supplementary Table 5). We aligned these data to the corresponding reference genomes and searched for splice junction reads whose 3’ splice sites mapped to positions orthologous to the identified *D. melanogaster* ratchet points. Despite having 2 orders of magnitude fewer reads from these species, 131 of the 197 *D. melanogaster* ratchet points (66.5%) were identified in at least one of the five other *Drosophila* species, 69 of which were identified in at least two species (Supplementary Table 6 & 7). Together these observations demonstrate that the nucleotide sequence and function of ratchet points are conserved among *Drosophila* species indicating that recursive splicing is evolutionarily conserved.

The host genes containing recursively spliced introns are expressed in a broad spectrum of developmental timepoints, tissues, and cell types – the recursive host genes are expressed at FPKM > 1 in 72%, 93% and 83% of cell lines, developmental time points, and tissues, respectively. However, host gene expression levels are quite dynamic throughout development and 63% have their peak expression in nervous system tissues (Fig. 4a), consistent with GO enrichments in development and neural functions (Supplementary Table 8).

**Figure 4.**
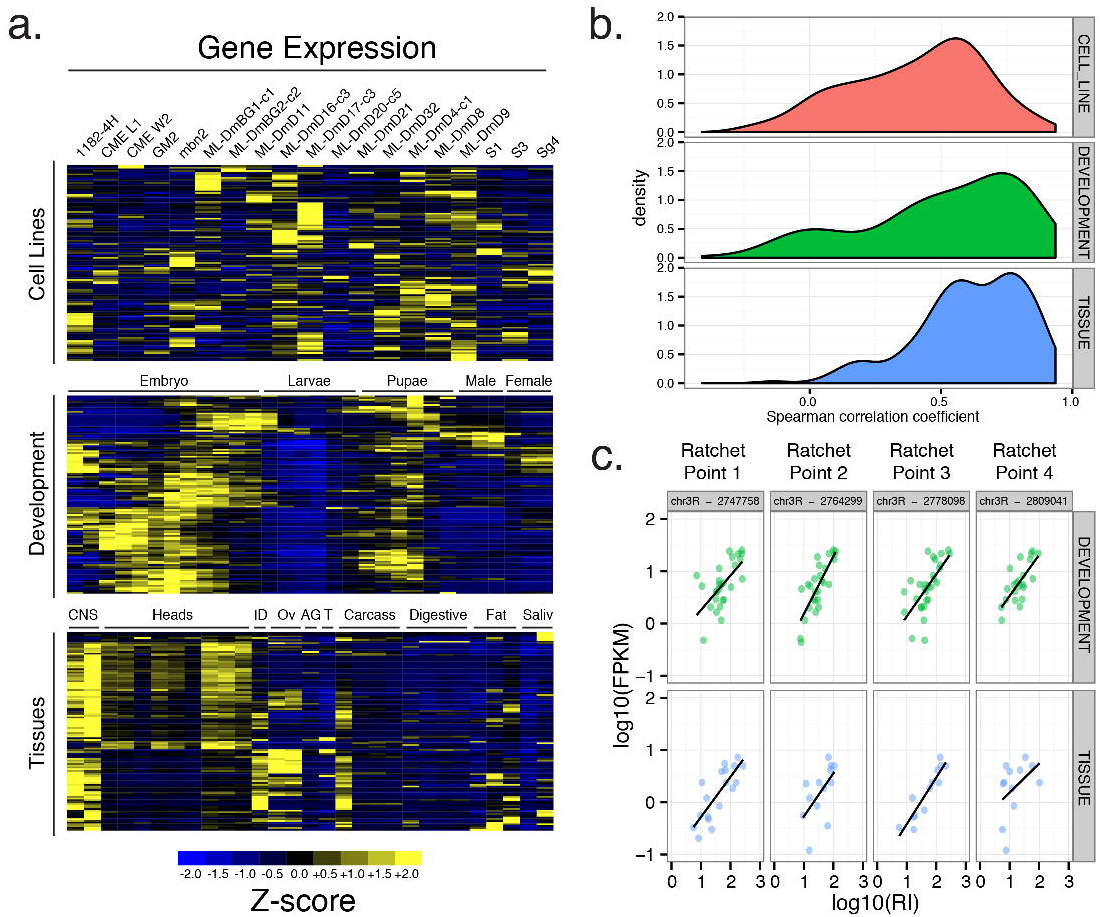
Expression characteristics of recursively spliced genes. **a,** Heatmap representation of Z-scores of mRNA expression levels of the recursively spliced genes among the samples examined. Imaginal discs (ID), ovaries (OV), accessory gland (AG) testes (T). **b,** Distribution of the Spearman correlations of mRNA expression levels and recursive indexes of each ratchet point for the cell line (red), developmental (green), and tissue (blue) samples. **c,** Example of the correlation of mRNA expression levels and recursive indexes for four ratchet points in *Antp* in the developmental (green), and tissue (blue) samples.

Importantly, in our analysis of lariat introns, although we considered all permutations of splicing events, we did not identify any reads corresponding to lariat introns that would be generated by direct splicing of the flanking constitutive exons without recursive splicing. This indicates that recursive splicing may be constitutive. To further assess this, we calculated a recursive index for each ratchet point (the number of ratchet point junction reads/mapped reads). We observed generally strong correlations between the recursive index and the gene expression level for most genes (Fig. 4b). For example, there is a strong positive correlation between gene expression and recursive splicing for all four ratchet points in the *Antennapedia* (*Antp)* gene (Fig. 4c). The correlation between gene expression and recursive splicing is strongest among the tissue samples and weakest among the cell lines, which have the highest and lowest number of mapped reads, respectively (Supplementary Fig. 4), indicating that low correlation is related to sequencing depth. Together, these results strongly suggest that recursive splicing is constitutive – when the host gene is transcribed, it is recursively spliced.

Recent studies have demonstrated strong associations between chromatin marks and particular features of gene architecture, including intron-exon boundaries. Of particular note, H3K4me3^(ref. 9)^, H3K79me2^(ref. 10)^, and H3K36me3^(ref. 11)^ have been shown to specifically transition near intron-exon boundaries in humans. We inspected ChIP-seq data obtained from whole larvae to determine whether any chromatin marks are associated with ratchet points (Supplementary Figure 5). None of the chromatin marks we examined are specifically associated with ratchet points, yet the recursive splice sites are associated with chromatin marks that would be expected given their position relative to canonical exons.

Here we provide experimental evidence that 130 *Drosophila* introns, 26 times the number previously known, are removed in multiple, sequential steps by recursive splicing, rather than by a single splicing event. The ratchet points involved in recursive splicing are highly conserved and share sequence similarity with one another. While recursive splicing clearly occurs, its function and mechanism remains elusive. As we have been unable to identify any features that differentiate recursive and non-recursive introns, it remains unknown why some introns are recursively spliced and others are not. Further investigation will be necessary to determine whether recursive splicing required for the function of the host gene and how the upstream exons re-engage in subsequent splicing reactions.

## METHODS SUMMARY

Total RNA strand-specific sequence data was generated using the Illumina TruSeq Total RNA Stranded kits and sequenced on an Illumina HiSeq 2000 using paired-end chemistry and 100-bp cycles. Sequences are available from the Short Read Archive (Accession numbers available in Supplementary Table 1). The total RNA-seq data was aligned using TopHat^12^. Recursive junction reads were independently identified from the TopHat alignments and from alignments to all possible recursive junction splice sites. The recursive splice sites were validated by RT-PCR of *Drosophila* S2 cell RNA and Sanger sequencing. Sequence logos were generated with WebLogo^13^. GO analysis was performed using Funcassociate 2.0^(ref. 14)^.

## Acknowledgments

This work was supported by NHGRI grant 1U54HG006994 to S.E.C. (PI) and B.R.G. (co-PI) as part of the modENCODE project.

## Author Contributions

B.R.G and S.E.C. supervised data production. S.O., A.O., and M.B. performed experiments. M.D., X.W., A.P., M.B., and B.R.G. performed computational analysis. B.R.G. wrote the paper with input from all authors.

## Author Information

Sequences are available from the Short Read Archive and the modENCODE website, a list of accession numbers is given in Supplementary Table 1. Reprints and permissions information is available at www.nature.com/reprints. The authors declare no competing financial interests. Readers are welcome to comment on the online version of the paper. Correspondence and requests for materials should be addressed to B.R.G.

